# The nucleoid as a scaffold for the assembly of bacterial signaling complexes

**DOI:** 10.1101/169912

**Authors:** Audrey Moine, Leon Espinosa, Eugenie Martineau, Mutum Yaikhomba, P J Jazleena, Deborah Byrne, Emanuele G. Biondi, Eugenio Notomista, Matteo Brilli, Virginie Molle, Pananghat Gayathri, Tâm Mignot, Emilia M.F. Mauriello

## Abstract

The FrzCD chemoreceptor from the gliding bacterium *Myxococcus xanthus* forms cytoplasmic clusters that occupy a large central region of the cell body also occupied by the nucleoid. In this work, we show that FrzCD directly binds to the nucleoid with its N-terminal positively charged tail and recruits active signaling complexes at this location. The FrzCD binding to the nucleoid occur in a DNA-sequence independent manner and leads to the formation of multiple distributed clusters that explore constrained areas. This organization might be required for cooperative interactions between clustered receptors as observed in membrane-bound chemosensory arrays.

**AUTHOR SUMMARY:** In this work, we show that the cytoplasmic chemoreceptor of the Frz chemosensory system, FrzCD, does not bind the cytoplasmic membrane like most MCPs but bind the bacterial nucleoid directly, thus forming distributed protein clusters also containing the Frz kinase. *In vitro* and *in vivo* experiments show that DNA-binding is not sequence-specific and is mediated by a basic aminoacid sequence of the FrzCD N-terminal domain. The deletion of this motif abolishes FrzCD DNA-binding and cooperativity in the response to signals. This work shows the importance of the nucleoid in the organization and functioning of cytoplasmic signaling systems in bacteria.

## INTRODUCTION

The bacterial cytoplasm is not a homogeneous solution of macromolecules, but rather a highly organized and compartmentalized space where the clustering and segregation of macromolecular complexes at certain cell regions confers functional efficiency [1]. Bacterial chemoreceptors represent a versatile model system to study the subcellular localization of macromolecules, as they are present in many prokaryotes where they form highly ordered arrays that occupy different positions in cells. Chemoreceptors, also called Methyl-accepting Chemotaxis Proteins (MCP), are capable of transducing external signals to downstream signaling pathways where phospho-cascades, initiating at the level of histine kinases CheAs and culminating at the level of output response regulators CheYs, translate the initial signal into cell behaviors such as regulation of motility, cell development or aggregation [2–4]. A common feature of the MCPs is their ability to form highly ordered hexagonal structures, which, by cryoelectron tomography, look like lattices with each unit composed of an MCP trimer of dimers, two CheW docking proteins and one CheA dimer [5–8]. Receptor clustering is not strictly required for signal transduction, as one functional unit is enough to generate phosphorylated CheY [6,9–11]. However, MCP clustering is essential to ensure the amplification of the initial signal, which is a direct consequence of the cooperative interactions between clustered chemoreceptors [11–15].

While the MCP lattices have been described for all studied prokaryotic chemoreceptors [16] their subcellular localization and distribution can vary among species, often reflecting life style complexity, behaviors and functions. For example, *Escherichia coli* MCPs localize in one or two polar clusters and more lateral clusters appear as cells become longer [17]. Differently, the TlpT cytoplasmic chemoreceptor from *Rhodobacter sphaeroides* forms a cluster positioned at the center of cells or two clusters positioned at the two and three cell quarters [18]. The determinants of these different localization patterns also vary. In *E. coli*, membrane-anchored MCPs form clusters stochastically and through a self-assembly mechanism [17]. The TlpA *Bacillus subtilis* polar chemoreceptor recognizes and associates with strongly curved membrane regions generated during cell septation. These regions become the new poles after cell division, which explains the TlpA polar localization [19]. While in *E. coli* and *B. subtilis* the polar targeting of bacterial chemoreceptors is due to intrinsic properties of these proteins, in *Vibrio* species the Che proteins are recruited to the cell poles by a set of specialized proteins responsible of the general maturation of these cell regions [20,21]. The presence of CheWs and CheAs also seem to be universally important in chemoreceptor cluster formation [22,23].

*Myxococcus xanthus* is a gliding bacterium that uses the Frz chemosensory system to modulate the frequency at which cells periodically reverse the direction of their movement on solid surfaces to reorient in the environment, analogously to controlled tumbles in *E. coli* [24]. The Frz regulation of directionality in *M. xanthus* is essential to achieve fruiting body formation, a behavior that bacteria initiate when they are exposed to unfavorable growth conditions. In the Frz pathway, the FrzCD chemoreceptor activates the autophosphorylation of a CheA-CheY fusion, FrzE, which in turn phosphorylates the response regulator FrzZ [25]. The system also possesses two CheW homologues (FrzA and FrzB), a methyltransferase (FrzF) and methylesterase (FrzG). The chemoreceptor of the Frz pathway, FrzCD, lacks the transmembrane and periplasmic domains, which are replaced by a N-terminal domain of unknown function [26]. When FrzCD was first localized in cells, it appeared organized in multiple dynamic cytoplasm clusters that aligned when cells made side-to-side contacts, which has been proposed to be part of a signaling process that synchronizes cell reversals [27]. However, the determinants of FrzCD localization and its exact link with the regulation of the cell reversal are still unclear.

In this work, we show that FrzCD forms cytoplasmic signaling clusters by directly binding to the nucleoid. This binding, is not absolutely required for signaling but it supports a cooperative dose-dependent response to external signals as detected in single cell assays. Thus, analogously to membrane chemotaxis clusters in *E. coli*, nucleoid binding could support the formation of chemosensory-like arrays with signal amplification properties.

## RESULTS

### The Frz chemosensory system co-localizes with the nucleoid of different bacterial species

Fluorescence microscopy has shown that a FrzCD-GFP fusion appears as multiple distributed clusters co-localizing with the nucleoid in *M. xanthus* cells (Figure 1A, 1B and Figure S1) [27–29]. It has been also shown that an inducible FrzE-YFP fusion colocalizes with FrzCD at the nucleoid [29]. To further analyze the co-localization of FrzCD and FrzE with the nucleoid, we first constructed a strain expressing a *frzE-mcherry* fusion that replaced the *frzE* locus and is, thus, expressed under its endogenous promoter. The chimeric protein was functional (Figure S2) and formed clusters very similar to that of FrzCD and also co-localizing with the *M. xanthus* nucleoid (Figure 1A, 1B and Figure S1).

**Figure 1.**
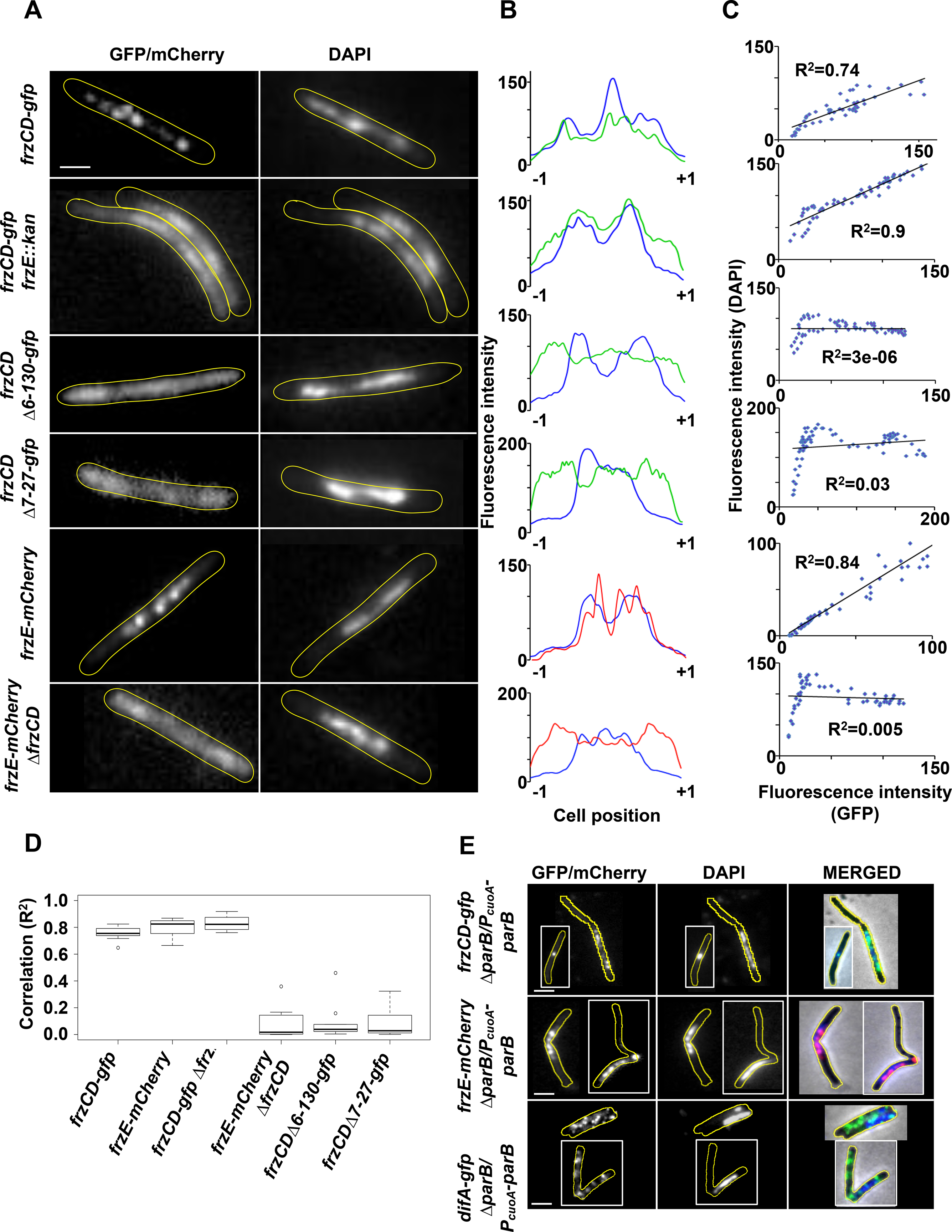
FrzCD-GFP colocalizes with the nucleoid in *M. xanthus*. **(A)** Micrographs of *M. xanthus* cells carrying a GFP or a mCherry fusion and stained with the DNA DAPI stain. The genetic backgrounds of the *M. xanthus* strains are indicated on the left. The white arrows indicate the cells whose fluorescence profiles and correlation coefficients between the DAPI and GFP localization are shown in (B) and (C), respectively. Cells surrounded by white boxes are taken from separate original micrographs. Scale bars correspond to 1μm. **(B)** GFP or mCherry (green or red) and DAPI fluorescence (blue) profiles with the fluorescence intensity (arbitrary units) represented on the *y* axis and the cell length positions with −1 and +1 indicating the poles, on the *x* axis. **(C)** Correlation coefficients between the DAPI and GFP or mCherry localization. R^2^ values > 0,5 indicate significant correlations. **(D)** Box plots indicate the medians of the correlation coefficients (R^2^) from 10 cells (from one biological replicate) of each of the indicated strains. **(E)** Micrographs of a *M. xanthus parB* conditional mutant carrying *frzCD-gfp* or *frzE-mcherry* and DAPI stained. Micrographs were obtained upon 18h depletion of ParB. The genetic backgrounds of the *M. xanthus* strains are indicated on the left. Scale bars correspond to 1μm.

To directly show that the nucleoid supports Frz protein localization, we constructed a *M. xanthus* conditional mutant that lacked ParB, a protein important for nucleoid segregation whose absence causes the presence of anucleated cells, cells with abnormal nucleoid condensation and cells where the division septum is improperly positioned over the nucleoid (“guillotines”) [30,31]. When *frzCD-gfp* and *frzE-mcherry* were expressed in the *parB* mutant, we observed that both FrzCD and FrzE clusters always co-localized with the nucleoid and cells lacking the nucleoid also lacked Frz clusters. Similarly, in cells with “guillotines”, septa formed in regions occupied by both the nucleoid and Frz clusters instead than in DNA-free regions (Figure 1E).

As a control, we looked at the cellular localization of another chemoreceptor fusion, DifA-GFP, in the absence of *parB*. DifA has been recently shown to form membrane bound and uniformly distributed clusters [28]. In the absence of *parB*, even cells without nucleoid still carried DifA-GFP clusters and these clusters localized similar to the wild type [28] (Figure 1E). These results confirmed that nucleoid-mediated cluster formation is a specific feature of Frz proteins.

To test if FrzCD and FrzE were capable of associating with the nucleoid independently of each other, we expressed *frzE-mcherry* in a strain lacking *frzCD* and *frzCD-gfp* in a strain lacking *frzE* (Mauriello et al., 2009 and this study). As previously shown, in the absence of FrzCD, FrzE-mCherry was homogeneously dispersed in the cytoplasm, and notably also in the polar regions (Figure 1A, 1B and Figure S1) [27,29]. Here we additionally showed that in the absence of *frzE*, FrzCD lost its typical punctate pattern, but the fluorescent signal was still retained towards the center of the cell body and strictly co-localized with the nucleoid (Figure 1A-D and Figure S1). The aberrant localization patterns observed in the deletion mutants were not due to a change in protein levels (Figure S3). Thus, this result confirms that both FrzCD and FrzE are important for cluster formation, whereas FrzCD is responsible for the recruitment of FrzE to the nucleoid.

To check whether the association between FrzCD and the nucleoid was *M. xanthus*-specific, we constructed an *E. coli* strain expressing *frzCD-gfp* from a plasmid and under the control of an IPTG inducible promoter. Under these conditions, FrzCD also co-localized with the nucleoid and this co-localization was particularly evident in elongated undivided cells containing multiple segregated nucleoids (Figure 2A-D).

**Figure 2.**
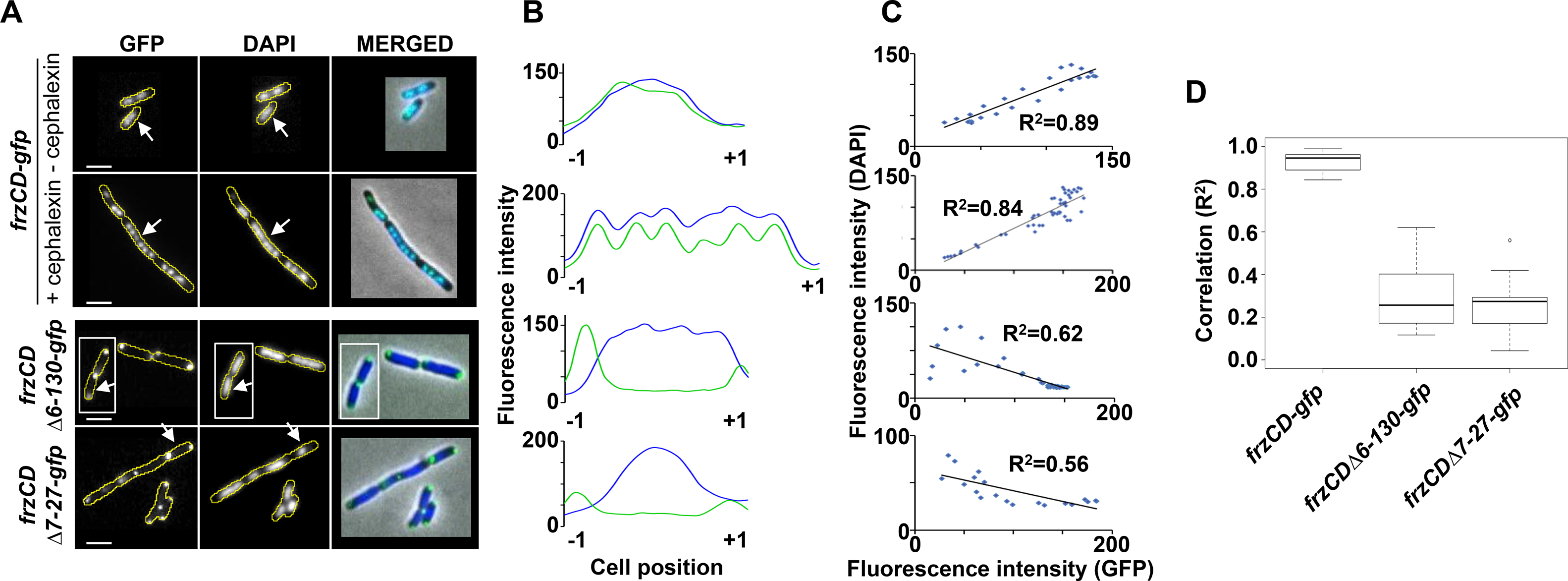
FrzCD-GFP colocalizes with the nucleoid in *E. coli*. **(A)** Micrographs of *E.coli* cells carrying a GFP fusion on a plasmid and stained with the DNA DAPI stain. The genetic fusions are indicated on the left. Cells carrying *frzCD-gfp* were also treated with 10μg/ml cephalexin to visualize FrzCD-GFP colocalization with the multiple nucleoids of undivided cells. The white arrows indicate the cells whose fluorescence profiles are shown in (B). Cells surrounded by white boxes are taken from separate original micrographs. Scale bars correspond to 1μm. **(B)** GFP (green) and DAPI fluorescence (blue) profiles with the fluorescence intensity (arbitrary units) represented on the *y* axis and the cell length positions with −1 and +1 indicating the poles, on the *x* axis. **(C)** Correlation coefficients between the DAPI and GFP localization. R^2^ values > 0,5 indicate significant correlations. **(D)** Box plots indicate the medians of the correlation coefficients (R^2^) from 10 cells (from one biological replicate) of each of the indicated strains.

These observations suggest that FrzCD can associate with the chromosomes of different bacterial species, either directly or by the aid of a docking factor common to *M. xanthus* and *E. coli*.

### FrzCD directly binds to the DNA

The possibility that FrzCD interacted with the nucleoid was puzzling especially considering that the direct binding between a chemoreceptor and the DNA has not been reported prior to this study. To explore this possibility, we generated a 6His-tagged FrzCD version, purified it from *E. coli* and tested its ability to form complexes with DNA. FrzCD binds directly to DNA because its presence altered DNA mobility on agarose gels (Figure 3A). Binding did not require extended DNA fragments because it was also observed with oligomers of DNA as small as 69 bp (Figure S4). Moreover, FrzCD DNA-binding did not seem to depend on the DNA sequence nor on its GC content (Figure S4) as anticipated by the *in vivo* results showing that FrzCD is distributed all over the nucleoid in the absence of FrzE (Figure 1A and B).

**Figure 3.**
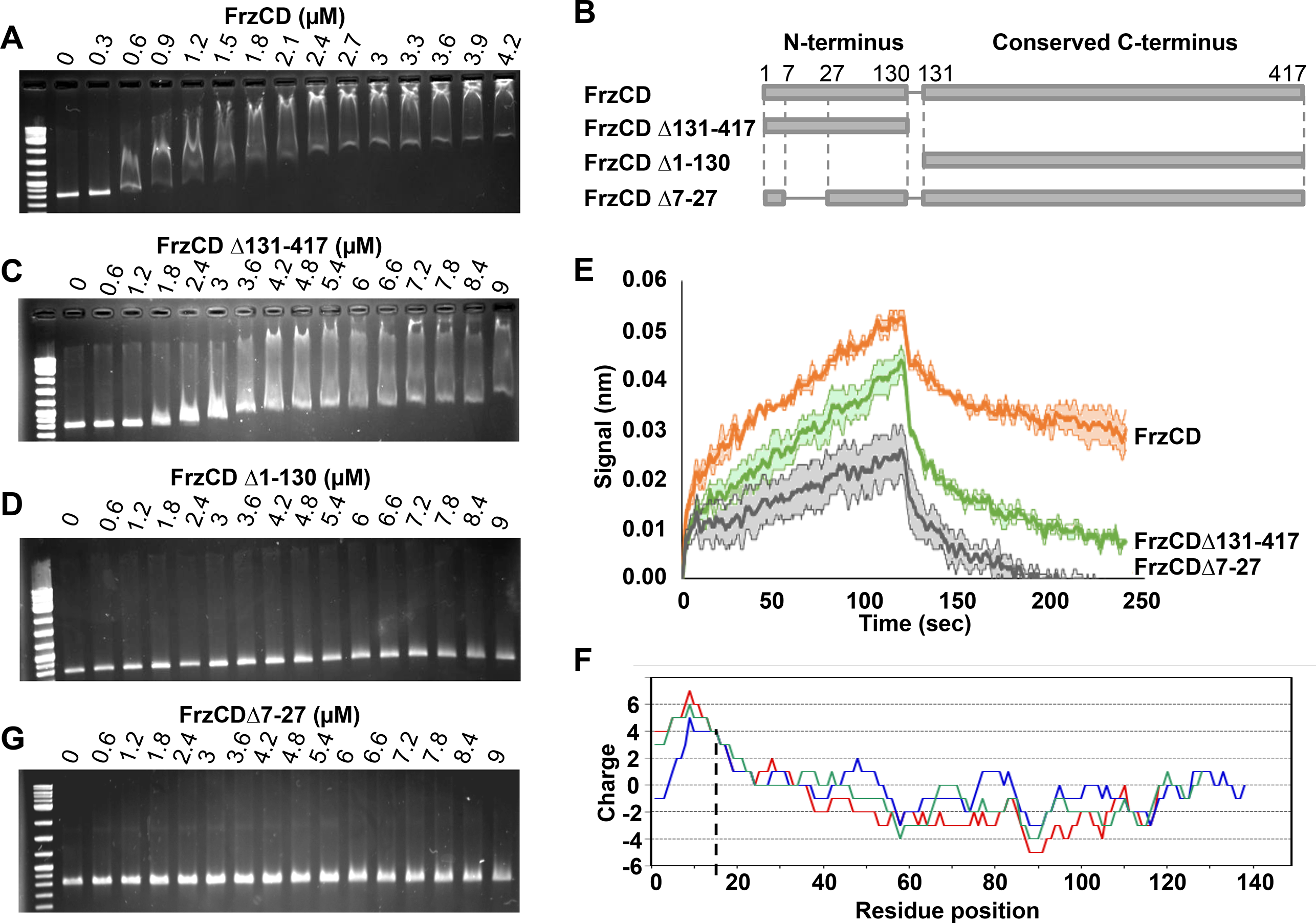
FrzCD directly interacts with the DNA *in vitro*. **(A)** Electrophoretic mobility shift assays (EMSA) on 1% agarose gels stained with ethidium bromide and developed at the UV light. The indicated concentrations of purified 6His-FrzCD were incubated with a 801 bp DNA fragment. **(B)** Schematic representation of the FrzCD protein domains. **(C-D)** The indicated increasing concentrations of 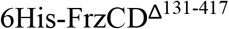 (C) and 6His-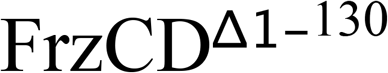 (D) were used in EMSA assays with a 801 bp DNA fragment. **(E)** Average binding curves and duplicates in degraded colors of each immobilized FrzCD construct 6His- FrzCD, 6His- 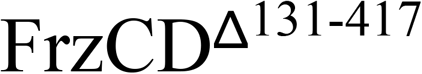 or 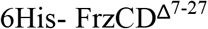, with a 474 bp DNA fragment at a concentration of 38nM. **(F)** “Sliding window” representation indicating the protein charge of the first FrzCD N-terminal region at the different positions and obtained with 10, 20 and 30 residue windows (blu, green and red, respectively). **(G)** Increasing concentrations of 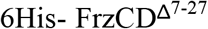 were used in EMSA assays. On the first lane of each gel, 500 ng of the 2-Log DNA ladder (0.1-10 kb, NEB) have been loaded. Data in panel (A, C, D, and G) are representative of three independent experiments.

The shift pattern depended on the FrzCD concentration (Figure 3A). More specifically, the shift of DNA fragments gradually increased as the FrzCD concentration was increased (Figure 3A and Figure S4). Such migration profiles have been previously described for proteins that can nucleate on DNA molecules in a non-specific manner, i.e. some Type Ib ParA-like proteins [32,33]. However, to exclude that such profiles were due to the formation of FrzCD unfolded aggregates, we checked the oligomerization state of our purified 6His-FrzCD. As expected for an MCP, FrzCD forms a homogenous dimer in solution, corresponding to a molecular weight of ~ 90 kDa (Figure S4). Last, 6His-FrzCD activated the autophosphorylation of the FrzE kinase *in vitro*, proving that it is functional (Figure S4).

While FrzCD does not appear to bind specific DNA motifs in vitro, it could bind to specific sites *in vivo* (perhaps with the help of additional factors), explaining the formation of clusters. To test this possibility, we performed chromatin-immunoprecipitation (ChIP) experiments using the *frzCD-gfp* strain and polyclonal GFP antibodies. As expected, FrzCD-GFP but not a FrzCD variant that cannot bind DNA (see below) was able to co-immunoprepicipate significant amounts of DNA. Deep-sequence (ChIP-Seq) [34] of the immunoprecipitated DNA revealed no enrichment in the pool of DNA fragments obtained with the ChIP meaning that FrzCD-GFP can bind any DNA sequence from the *M. xanthus* genome (Figure S5; the ChipSeq results have also been deposited on GEO (https://www.ncbi.nlm.nih.gov/geo/, accession number GSE102724) and an accession number is being created). As a positive control we used a *parB-yfp* strain [30]. In this case, as expected, the nucleoid region corresponding to positions 9,109 to 9,110 Kb and containing *parS* [30] was highly represented in the DNA pool obtained with ParB-YFP (Figure S5). We conclude that FrzCD binds DNA in a non-sequence-specific manner to recruit FrzE and thus form clusters containing Frz signaling complexes.

### The FrzCD N-terminal region is required for the FrzCD DNA binding

Beside a very conserved C-terminal methylation domain, FrzCD contains a unique 137 residue N-terminal region (Figure 3B). We then asked whether this region corresponded to the FrzCD nucleoid-binding domain and tested its ability to form complexes with the DNA in our gel shift assays. Indeed, the FrzCD N-terminal domain (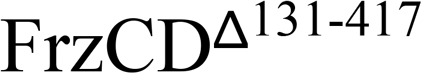) also provoked a mobility shift of DNA fragments of different length, albeit at a lower efficiency (compare Figure 3A and C). On the other hand, the FrzCD C-terminal methyl-accepting domain alone (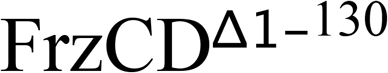) did not associate with any DNA fragment (Figure 3D) showing that the methylation domain does not bind DNA.

To confirm a direct interaction between FrzCD and DNA and also better compare the DNA-binding properties of FrzCD and 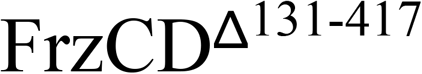, we performed DNA-protein interaction experiments using Bio-Layer Interferometry (BLI), a technique previously used to study protein-protein interactions [35]. FrzCD and DNA interaction was also detected in this assay (Figure 3E). Consistent with the gel-shift experiments, binding appeared complex and could not be fitted to a 1:1 interaction model, precluding precise determination of a K_D_. Nevertheless, when we compared the DNA binding curves of 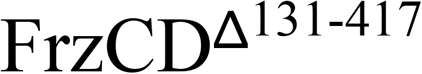 and FrzCD, the results confirmed that 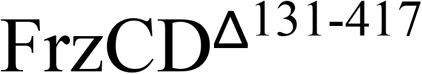 binds the DNA at a lower efficiency than FrzCD (showing slower association and faster dissociation, Figure 3E).

By further analyzing the FrzCD N-terminal domain, we realized that it contains a positively charged peptide of approximately 30 amino acids followed by a more negative region predicted to contain alpha helices (Figure 3B and F, Figure S6). By searching for homologs in the UniProtKB/SwissProt Data Base, we found that such FrzCD N-terminal basic peptide was similar to the basic tail present at the N terminus of eukaryotic histones (Figure S6) whose deletion has been shown to substantially affect histone-DNA interactions and decrease nucleosome stability [36,37]. To test whether this sequence had a histone-tail-like function in the binding of FrzCD to DNA *in vitro*, we generated a 6His-tagged FrzCD version only lacking the basic amino acid sequence from residue 7 to 27 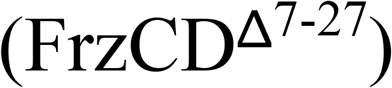, purified it from *E. coli* and tested its ability to form complexes with the DNA in our gel shift assays. Remarkably, 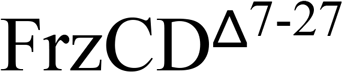 did not shift the migration of DNA fragments on agarose gels, similarly to 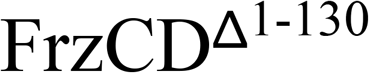 missing the entire N-terminal domain (Figure 3D and 3G). In the BLI assay, binding of 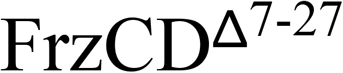 to DNA was still detectable, however it was severely impaired (Figure 3E). This result suggests that the positively charged motif is required for efficient DNA binding but it may not be the sole determinant.

The different DNA binding efficiencies of the four FrzCD, 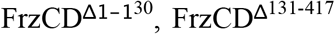 and 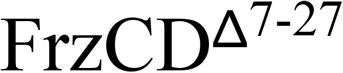 alleles were not due to altered protein stability and folding because all recombinant proteins, except as expected 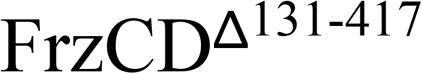 (the signaling domain) were able to activate the autophosphorylation of the FrzE kinase *in vitro* (Figure S4).

### The binding of FrzCD to the nucleoid is required for FrzCD cluster formation *in vivo*

To check whether the absence of the N terminus or the basic tail also affected the binding of FrzCD to DNA *in vivo*, we used *M. xanthus* strains expressing 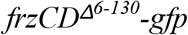 or 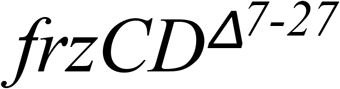-*gfp* at the *frzCD* locus (Mauriello *et al.*, 2009 and this study). The 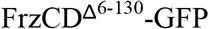 and 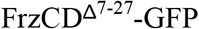 fluorescence appeared mostly diffused, also occupying the polar regions (Figure 1A-D). The two protein fusions could only rarely form short-lived clusters that localized anywhere in the cells (not only at the central region). In all cases, the aberrant localization patterns were not due to protein stability (Figure S3). In addition, when 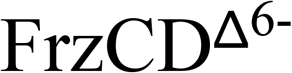 ^130^-GFP and 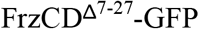 were produced in *E. coli* they also lost their co-localization with the nucleoid. However, instead of looking dispersed in the cytoplasm as in *M. xanthus*, 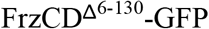 and 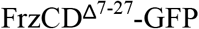 were confined at one cell pole in *E. coli*. It is likely that 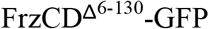 and 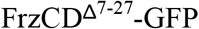 formed aggregates targeted to the poles due to the absence of the nucleoid anchor (Figure 2).

### FrzCD cluster dynamics and organization

To understand how FrzCD clusters are formed along the nucleoid, we analyzed the cluster dynamics at high temporal resolution. Contrarily to previous assumptions based on lower resolution analysis [27], this analysis showed that FrzCD clusters are quite fixed and only featured by Brownian-like motions in highly constrained areas of the nucleoid (Figure 4A-C). This mobility decreased with the increase of cluster intensity, suggesting that clusters containing more molecules might be more tightly anchored to the chromosome and, hence, more fixed (Figure 4D). To test whether the signaling state of FrzCD also affects the cluster mobility, we tested whether clusters were also constrained in strains carrying point mutations in methylation sites either generating FrzCD loss of function (*frzCD*_*E202A-E203A*_*::gfp*) or, oppositely, FrzCD gain of function (*frzCD*_*E168A-G169A*_*::gfp*) [25,27,39]. There were no notable differences between the tested conditions, suggesting that signaling does not affect the nucleoid dynamics of Frz signaling complexes along the nucleoid (Figure 4E).

**Figure 4.**
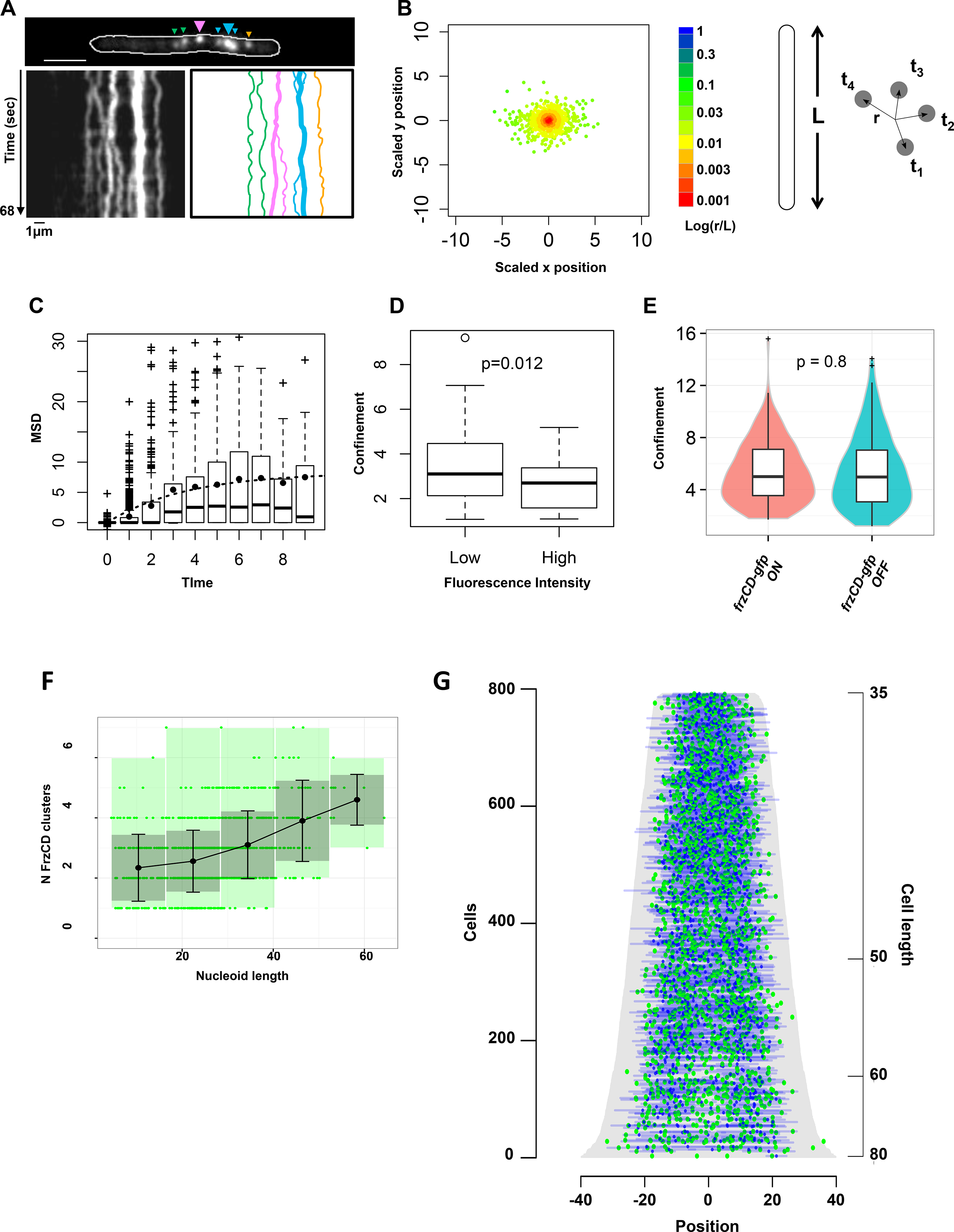
The organization of FrzCD clusters depends on cluster intensity and mobility. **(A)** A representative fluorescence 1 second time-lapse (left panel) and the corresponding kymograph (middle panel) of a *frzCD-gfp* cell (top panel). Big and small arrows indicate large and small clusters, respectively. The right panel represents the trajectories of each cluster (same color codes as on the top panel). Scale bars correspond to 1μm. **(B)** Cluster displacement (r) from the mean position at each given time (t). L represents the cell length. The color code corresponding to the logarithm of the ratio r/L indicates that the amplitude of the cluster displacement never exceeds 5% of the cell length. **(C)** Box plots indicate the distribution of the Mean Square Displacements at the different lag times; the mean of each lag value is indicate by the black dots. **(D)** Box plots indicate a significant decrease of the median confinement for clusters of low fluorescence intensity compared to high intensity clusters. For panels B, C and D 1039 clusters from 297 cells (two biological replicates) were analyzed. **(E)** The box plots and the violin plots show the measured confinements of *frzCD-gfp* strains blocked in the ON (*frzCD*_*E168A-G169A*_*::gfp*, hyper-reversing) and OFF (*frzCD*_*E202A-E203A*_*::gfp*, hypo-reversing) states. 130 and 150 clusters were analyzed for the ON and OFF states, respectively. **(F)**. Average numbers of clusters with standard deviations (black dots and bars, respectively) for different nucleoid sizes are shown. Green dots represent measurements for individual cells. Grey zones represent the variances. 2564 clusters from two biological replicates were analyzed. **(G)** 909 cells were ordered according to their cell length (pixels, grey) and for each cells GFP and DAPI fluorescence are represented as green and blue dots, respectively, at their corresponding cell position. 0 is the cell center.

Given that FrzCD clusters are not very dynamic and form in a limited number of sites per chromosome, we wondered how these clusters were segregated to daughter cells following cell division. Analysis of the number of FrzCD clusters as a function of chromosome length revealed a linear correlation (Figure 4F), suggesting that more clusters are assembled as the DNA is being replicated. A demograph representation revealed that the clusters are statistically distributed along replicating as well as segregated chromosomes (Figure 4G). This distribution might ensure equal partitioning in daughter cells.

### The nucleoid-dependent assembly is important for cooperative signal-dependent response

Similar to transmembrane chemosensory clusters [6,9–11], the formation of Frz nucleoid-associated clusters is not strictly required for signaling. In fact, it has been previously shown that a 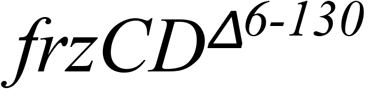 strain, where FrzCD molecules can no longer bind DNA and are, thus, diffused (Figure 1) can still produce reversals (Bustamante et al., 2004). In *E. coli* chemosensory cluster formation is also not critical for signal transduction but it confers properties such as the amplification of signal, a direct consequence of the cooperative interactions between clustered chemoreceptors [11–15]. Thus, we decided to check if the formation of DNA-bound Frz clusters can also promote cooperativity in the signaling activity of the Frz chemosensory system. For this, we took advantage of a newly developed microfluidic single cell assay where the frequency of reversals can be measured as a function of increasing concentrations of an artificial Frz-signal activator, isoamyl alcohol (IAA) [25]. Consistent with previous observations [25], in cells where FrzCD formed nucleoid bound clusters, IAA induced a dose-dependent response with a sigmoidal shaped curve that could be fitted by the Hill equation with a coefficient *n* = 3.017 +/− 0.2 (P = 0.0007), thus revealing possible cooperativity in IAA-dependent Frz activation (Figure 5A). Such response is FrzCD-dependent because a *ΔfrzCD* strain does not reverse at any IAA dose (Figure 5A). In the 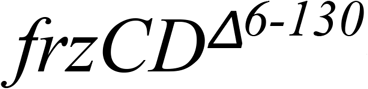 mutant, the cells still responded to IAA in a dose-dependent manner but in this case, the calculated Hill coefficient was *n* = 1.15 +/− 0.01 (P = 0.008), revealing that cooperation is lost in this mutant. These results suggest FrzCD clusters have signal amplification properties that are absent in the 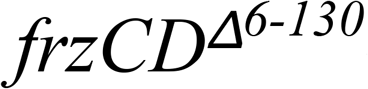 mutant. This property is advantageous to *M. xanthus* social behaviors because swarming and predation are defective (although moderately) in the 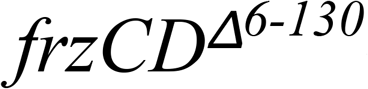 mutant compared to the wild type (Figure 5B and C).

**Figure 5.**
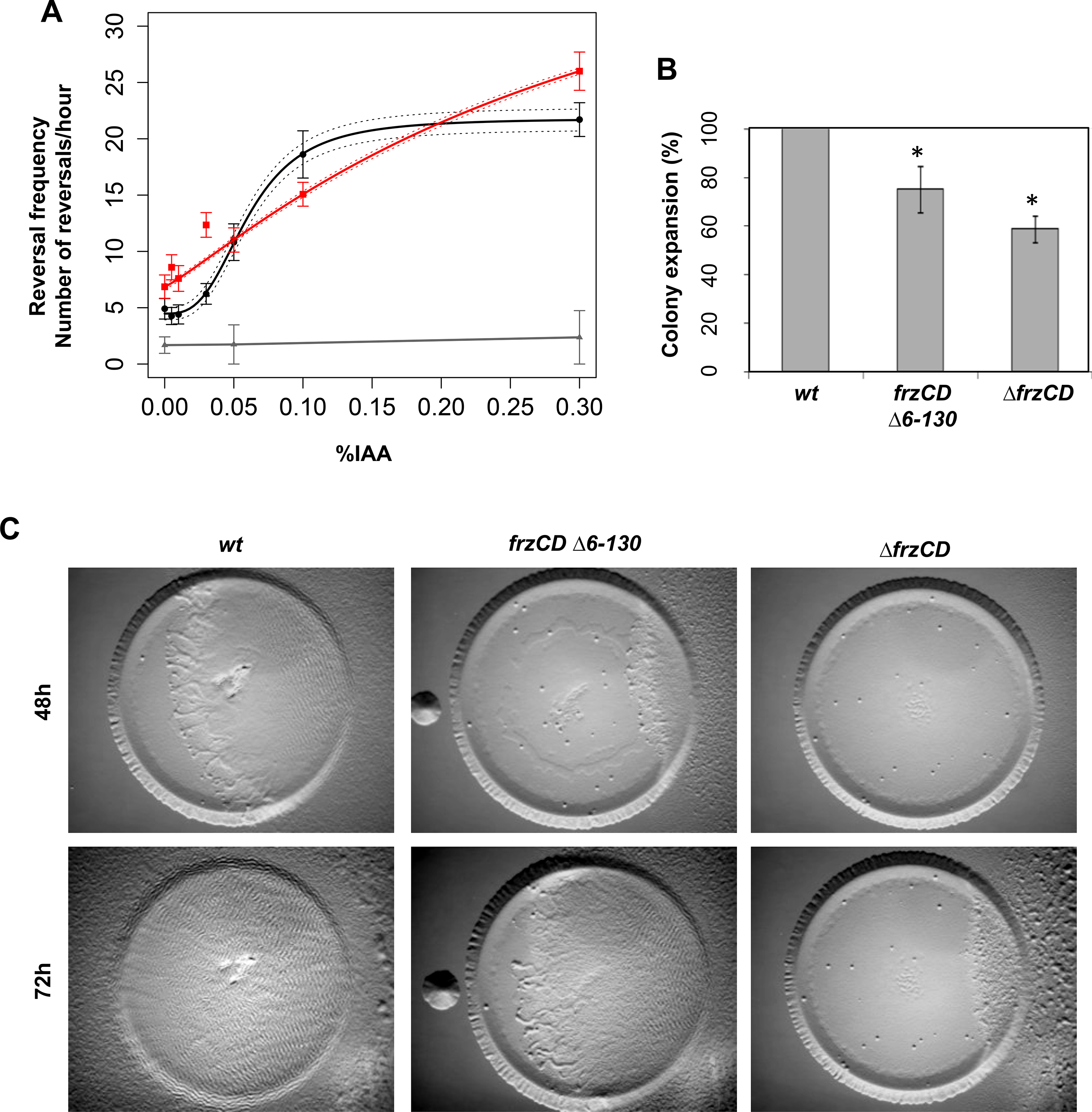
Frz cluster formation generates signal sensitivity in turn important for social behaviors. **(A)** The average reversal frequencies, calculated by scoring FrzS-YFP pole-to-pole oscillations are shown as a function of the IAA concentration for wild type (black), *frzCD*^*Δ6-130*^ (red) and *ΔfrzCD* (grey). Reversal frequencies values of wild type and *frzCD*^*Δ6-130*^ can be fitted by the Hill equation with an interval of confidence of 95% (dashed lines). Error bars represent the standard errors of the means. Reversal frequency values for each IAA dose and each strain are the results of a biological duplicate (each being a technical triplicate). About one hundred cells for the wild type and *frzCD*^*Δ1-130*^ strains and fifty for the *ΔfrzCD* strain were analyzed (refer to Supplementary Table 3 for the exact number of analyzed cells for each strain and IAA doses used in this experiment). **(B)** Colony expansion of wild type, *frzCD*^*Δ6-130*^ and *ΔfrzCD* cells. Error bars represent the standard deviations of the means from three biological replicates. **(C)** The same strains were analyzed in *E. coli* predation assays.

## DISCUSSION

In bacteria, transmembrane chemoreceptors use the bacterial inner membrane as a platform to form the well described arrays of trimers of dimers [41]. Remarkably, the *M. xanthus* Frz system forms clusters not on the membrane, but directly on the bacterial chromosome. Cluster assembly is directed by the chemoreceptor FrzCD, which binds to the DNA by a N-terminal domain carrying a positively charged tail similar to that found in eukaryotic histones [36,37]. While the binding of FrzCD to DNA is essential to target the Frz chemosensory system to the nucleoid, it is not sufficient for Frz cluster formation, as it requires downstream interactions with the FrzE kinase. Because FrzCD appears to bind DNA in a non-sequence specific manner, DNA-bound clusters do not occupy fixed localization sites and move across small areas on the nucleoid surface. Analogous to trans-membrane proteins diffusing in the bacterial membrane, the FrzCD cluster dynamic behavior may be affected by the size of the complex (and thus the number of interactions with DNA), explaining why bright clusters show Brownian-like motions that only explore constrained nucleoid areas.

Several lines of evidence suggest that the Frz cluster formation on the nucleoid occurs in a stochastic manner similarly to the assembly of the *E. coli* Che lattices in the membrane. First, the initial binding of FrzCD to DNA might take place anywhere on the nucleoid as such binding is not DNA-sequence specific. Once recruited to the nucleoid, small FrzCD foci diffuse, non-directionally, across confined small areas until they might nucleate large fixed clusters by attracting more FrzCD molecules. “Newborn” FrzCD foci might also, at one point, be incorporated by existing neighboring clusters. Thus, the areas explored by FrzCD clusters might represent the minimal critical distance from other clusters at which foci can exist. The existence of such minimal critical distance is supported by the fact that the number of Frz clusters increases linearly with the nucleoid size, suggesting that more clusters can form when more surface becomes available. Thus, like for transmembrane chemoreceptors, FrzCD molecules might either nucleate new dynamic foci if they are far enough from existing clusters, or encounter and join neighboring clusters. This simple mechanism might ensure correct partitioning of the clusters in daughter cells in the absence of further active mechanisms.

Cluster formation is unlikely to be uniquely linked to partitioning constraints during cell division. Indeed, the FrzCD variants that do not bind to the nucleoid form diffused signaling complexes that can be easily partitioned between daughter cells via diffusion. We therefore favor that nucleoid binding confers particular signaling properties to Frz complexes. In support of this, chromosome-bound complexes respond to signals in a cooperative manner, while soluble complexes do not. This could be explained if Frz clusters were arranged on the chromosome similar to the transmembrane chemoreceptors [42], forming ordered signaling arrays with signal amplification properties.

But, what signals are sensed by the chromosomal complexes? Frz clusters have been previously associated to the regulation of the cell reversal frequency in response to cell-cell contact. In fact, FrzCD clusters align in adjacent*M. xanthus* cells, a behavior that also seemed to induce cell reversals [27]. In support of this hypothesis, adjacent cells of a *frzE* strain did not produce cluster alignment. We now find that what seemed to be more diffused FrzCD clusters in the *frzE* strain at the time, are, in fact, FrzCD molecules dispersed on the nucleoid. Moreover, we now show that the Frz cluster dynamics are independent on the Frz signaling activity. Thus, the observed FrzCD cluster alignment might be more likely determined by similar dynamic rearrangements of the nucleoid of adjacent cells rather than by an active regulated mechanism. Therefore, we propose that chromosome assembly allows the formation of ordered chemosensory complexes that sense a yet unidentified intracellular signal. Finally, it cannot be excluded that the Frz complex responds to or even regulates chromosomal processes, although there is currently no evidence for such activity of the Frz system.

Finally, the analysis of the FrzCD sequences from some related species of δ-proteobacteria shows that while the FrzCD C-terminal region is very conserved, its N-terminus largely varies. Nevertheless, the FrzCD N-terminus always shows a positively charged sequence (Figure S7) suggesting that the non-sequence-specific recruitment of Frz proteins to the nucleoid essentially requires the presence of a positively charged protein domain rather than a specific amino acid sequence.

This type of cellular organization may be common to other bacterial macromolecular complexes to provide important regulatory functions. In this sense, the Frz example provides new perspectives to the role of the bacterial nucleoid as a scaffold for the spatial control of cellular functions.

## MATERIALS AND METHODS

### Bacterial strains, plasmids and growth

Strains and plasmids are listed in Table S1 and S2. *M. xanthus* strains were grown at 32°C in CYE rich media as previously described. *Pcuo::parB-ΔparB* cells were grown at 32°C in CTT supplied with 300 μM CuSO_4_.

Plasmids were introduced into *M. xanthus* cells by electroporation. Deletions and GFP fusions were inserted in frame to avoid polar effects on the downstream gene expression. These strains were obtained by homologous recombination based on a previously reported method using the pBJ113 or pBJ114 vectors [26,28]. To generate strains expressing GFP or mCherry fusion proteins, we constructed DNA cassettes including the last approximately 800 bp of each gene, with the exception of the stop codon; the gene encoding the *egfp* or *mcherry* genes from pEGFP-N1 (Invitrogen) or pEM147 [38] excluding the start codon and including the stop codon; the intergenic region between the gene of interest and its immediately downstream gene, if any; the first 800 bp of the downstream gene.

To construct the *parB* conditional mutant, we transferred the previously described *parB* conditional depletion [30] in our wild type strain, DZ2 [30,31].

To construct strains carrying a FrzS-YFP fusion, we used the pEFrzSY plasmid [25] *Escherichia coli* cells were grown under standard laboratory conditions in Luria-Bertani broth supplemented with antibiotics if necessary.

For swarming assays, cells (5 μl) at a concentration of 5×10^9^ cfu ml^−1^ were spotted on CYE agar plates and incubated at 32°C and photographed after respectively 48h with an Olympus SZ61 binocular stereoscope. For predation assays, *E. coli* (3μl at a concentration of 5×10^9^ cfu ml^−1^) and *M. xanthus* cells (3μl at a concentration of 5×10^9^ cfu ml^−1^) were spotted at the average 0.7 mm distance from each other on CF agar plates, incubated at 32°C and photographed after 72 h.

### Protein purification

BL21(DE3) [F^−^ ompT hsdSB(rB^−^ mB^−^) gal dcm (DE3)] cells were grown in Luria-Bertani broth supplemented 100 μg/ml ampicillin to mid-exponential phase at 37°C. For the experiments shown on Figure 3 and phosphorylation assays on Figure S4, overexpression was induced by adding 0,1 mM IPTG for cells containing plasmid pEM414 or 0,5mM for pEM415 and pEM433. Cells were then grown at 16°C over night. Cells were washed and resuspended in lysis buffer (50 mM TrisHCl, pH 8; 300 mM NaCl; 100 μg/ml PMSF; 30 U/mL Benzonase) and lysed at the French press. The cell lysates were centrifuged at 4°C for 30 min at 13000× rpm. Soluble tagged His_6_-proteins were purified on 1ml HisTrap^TM^FF columns (GE Healthcare) and desalted with PD-10 columns (GE Healthcare). Ultimately, purified proteins were eluted in 50 mM TrisHCl, pH 8 and 300 mM NaCl.

For the experiments shown on Figure S4A-E, the C-terminal-tagged FrzCD-(His)_6_ was expressed in *E. coli* BL21-AI (Invitrogen) cells. The cells were grown in Luria-Bertani broth containing 100 μg/ml ampicillin at 37°C till OD_600_ of 0.8 – 1.0, and incubated at 30°C for 5 hours post induction. The harvested cells were resuspended in Buffer A (50 mM Tris, 200 mM NaCl, pH 8.0) containing 10 % glycerol and lysed by sonication (Sonics VibraCell, 5 minutes, 60% amplitude, 1” ON, 3” OFF cycle). The cell lysate was spun at 39, 191 g for an hour and the supernatant loaded to a 5-ml HisTrap^TM^ FF (GE Healthcare) equilibrated with Buffer A. Following wash and elution with increasing concentrations (5%, 10%, 20%, 50% and 100%) of Buffer B (50 mM Tris, 200 mM NaCl, 500 mM imidazole, pH 8.0), pure fractions containing FrzCD-H_6_ were pooled and dialysed into buffer containing 50 mM Tris, 25 mM NaCl, 1 mM EDTA, pH 8.0 to enable further purification through ion exchange chromatography using a MonoQ-PE (GE Healthcare) column. The fractions containing the protein were pooled and concentrated using centricons. The protein was finally in buffer containing 50 mM NaCl, 1 mM EDTA and 50 mM Tris pH 8.0 either through dialysis or by buffer exchange during the concentration step. The concentrated protein was flash frozen and stored in small aliquots in −80°C till further use.

### Electrophoretic mobility shift assays (EMSA)

EMSAs on Figure 3 were carried out by incubating different concentrations of purified proteins with 10 nM PCR-amplification products of different sizes (Table S4), in buffer (10 mM of TrisHCl at pH 8; 60 mM of NaCl; 10% glycerol). Reactions were incubated for 40 min at 4°C before being loaded on 1% agarose gels. Gel migration was performed in 1X TBE at 4°C for 55 min. Gels were, then, stained with ethidium bromide and revealed at the UV light.

EMSAs for shorter oligos (Figure S4 and Table S4) were carried out in buffer containing 50 mM Tris, 50 mM NaCl, 1 mM dithiothreitol and 10 % glycerol, pH 7.4. All the oligonucleotide fragments were PCR-amplification products, except the 69 bp DNA that was generated by annealing the custom-synthesized oligonucleotides (Sigma) corresponding to the 5’ to 3’ sequence and its complementary strand (Table S4). The samples were incubated at 25°C for 20 min and loaded onto the appropriate % of agarose gel (1% for 1.3 kbp, and electrophoresed for 60 - 90 minutes in 1x TAE (40 mM Tris-Acetate and 1 mM EDTA).

### Oligomerization study using Size Exclusion Chromatography coupled with Multi-Angle light Scattering (SEC-MALS)

The expected elongated structure of an MCP precludes estimation of the oligomeric status by size exclusion chromatography alone, and hence we carried out SEC coupled with multi-angle light scattering to estimate the molecular mass in solution. The mass was determined using a Wyatt Dawn Heleos II equipped with light scattering detectors at 18 angles and an Optilab TrEX differential refractive index detector. Protein sample (100 μl of 2 mg/ml solution or 45 μM) was injected into the size exclusion column (BioRad EnRich650) equilibrated with the buffer 50 mM Tris, 50 mM NaCl, pH 8.0, and the run carried out at a flow rate of 0.4 ml/min. The observed molecular masses at various points along the peak in the elution curve were calculated using the protein concentration estimated from the differential refractive index (dRI), and measured scattered intensities. The Debye model in the ASTRA software provided with the equipment was used for fitting the data.

### Biolayer interferometry

Protein-DNA interaction experiments were conducted at 25°C with the BLItz instrument from ForteBio (Menlo Park, CA, USA). The BLI consists in a real time optical biosensing technique exploits the interference pattern of white light reflected from two surfaces to measure biomolecular interactions [45]. Purified 6His-FrzCD, 6His-FrzCD^Δ131-417^, 6His-FrzCD^Δ6-130^ and 6His-FrzCD^Δ7-27^ protein ligands were immobilized onto two different Ni-NTA biosensors (ForteBio) in duplicate at 1μM concentrations. A PCR amplified DNA fragment (474bp) with primers AGACCCCCGCACCCACGGAG and TCACGCGGGCTCCGGCTC (Eurogentec) was used as the analyte throughout the study at the 38nM. The assay was conducted in PBS pH 7.5, 0.001% tween-20. The binding reactions were performed with an initial baseline during 30 seconds, an association step at 120 seconds and a dissociation step of 120 seconds with lateral shaking at 2200rpm. A double reference subtraction (sensor reference and 6His-FrzCD^Δ1-130^) was applied to account for non-specific binding, background, and signal drift to minimize sensor variability.

### Chromatin Immunoprecipitation-deep sequencing (ChIp-seq)

ChIp-seq was performed as previously described [34]. In particular, mid-log phase cells (80 ml, OD_600_ of 0.6) were cross-linked in 10 mM sodium phosphate (pH 7.6) and 1% formaldehyde at room temperature for 10 min and on ice for 30 min thereafter, washed thrice in phosphate buffered saline (PBS) and lysed with lysozyme 2.2 mg ml^−1^ in TES (Tris-HCl 10 mM pH 7.5, EDTA 1 mM, NaCl 100 mM). Lysates (Final volume 1ml) were sonicated (Branson Digital Sonicator 450, Branson Sonic Power. Co., www.bransonic.com/) on ice using 10 bursts of 30 sec (50% duty) at 30% amplitude to shear DNA fragments to an average length of 0.3-0.5 kbp and cleared by centrifugation at 14,000 rpm for 2 min at 4°C. Lysates were normalized by protein content by measuring the absorbance at 280 nm; ca. 7.5 mg of protein was diluted in 1 mL of ChIP buffer (0.01% SDS, 1.1% Triton X-100, 1.2 mM EDTA, 16.7 mM Tris-HCl [pH 8.1], 167 mM NaCl plus protease inhibitors (Euromedex, https://www.euromedex.com/) and pre-cleared with 80 μL of protein-A agarose (Sigma-Aldrich, www.sigmaaldrich.com) and 100 μg BSA. Polyclonal GFP antibodies were added to the remains of the supernatant (1:1,000 dilution), incubated overnight at 4°C with 80 μL of protein-A agarose beads pre-saturated with BSA, washed once with low salt buffer (0.1% SDS, 1% Triton X-100, 2 mM EDTA, 20 mM Tris-HCl (pH 8.1), 150 mM NaCl), high salt buffer (0.1% SDS, 1% Triton X-100, 2 mM EDTA, 20 mM Tris-HCl (pH 8.1), 500 mM NaCl) and LiCl buffer (0.25 M LiCl, 1% NP-40, 1% sodium deoxycholate, 1 mM EDTA, 10 mM Tris-HCl (pH 8.1) and twice with TE buffer (10 mM Tris-HCl (pH 8.1) and 1 mM EDTA). The protein-DNA complexes were eluted in 500 μL freshly prepared elution buffer (1% SDS, 0.1 M NaHCO3), supplemented with NaCl to a final concentration of 300 mM and incubated overnight at 65°C to reverse the crosslinks. The samples were treated with 2 μg of Proteinase K for 2 h at 45°C in 40 mM EDTA and 40 mM Tris-HCl (pH 6.5). DNA was extracted using QIAgen minelute kit and resuspended in 30 μl of Elution Buffer. ChIp DNA sequencing was performed using Illumina MiSeq and analyzed using Galaxy Web Portal (usegalaxy.org/). Reads were analyzed by MatLab.

### Protein sequence analyses

In order to search for homologs of the FrzCD N-terminal domain, the first 130 aminoacids of the FrzCD sequence were BLAST into the UniProtKB/SwissProt Data Base (http://blast.ncbi.nlm.nih.gov). Predictions of secondary structures and protein sequence alignments were obtained with Jpred [46] and Clustal Omega [47], respectively. To analyze the FrzCD N-terminal region protein charge, “Sliding window” analyses were performed with Microsoft Excel.

### *In vitro* autophosphorylation assay

*In vitro* phosphorylation assays were performed with *E. coli* purified recombinant proteins. 4 μg of FrzE^kinase^ were incubated with 1μg of FrzA and increasing concentrations (3μg) of different FrzCD proteins (entire FrzCD, FrzCD^c^, FrzCDΔ6-130, FrzCDΔ7-27 or FrzCDΔ131- 417) in 25 μl of buffer P (50 mM Tris-HCl, pH 7.5; 1 mM DTT; 5 mM MgCh; 50mM KCl; 5 mM EDTA; 50μM ATP, 10% glycerol) supplemented with 200 μCi ml^−1^ (65 nM) of [γ-33P]ATP (PerkinElmer, 3000 Ci mmol^−1^) for 10 minutes at room temperature in order to obtain the optimal FrzE^kinase^ autophosphorylation activity. Each reaction mixture was stopped by addition of 5 × Laemmli and quickly loaded onto SDS-PAGE gel. After electrophoresis, proteins were revealed using Coomassie Brilliant Blue before gel drying. Radioactive proteins were visualized by autoradiography using direct exposure to film (Carestream).

### Fluorescence microscopy and image analysis

For fluorescence microscopy analyses, 5 μl of cells from 4 × 10^8^ cfu ml^−1^ vegetative CYE cultures were spotted on a thin fresh TPM agar pad at the top a slide (Mignot *et al.*, 2005). A cover slip was added immediately on the top of the pad, and the obtained slide was analyzed by microscopy using a Nikon Eclipse TE2000 E PFS inverted epifluorescence microscope (100 x oil objective NA 1.3 Phase Contrast) [48].

To study the colocalization with the DNA, the TPM agar pads were supplied with 1μg/ml DAPI stain and 1 mM IPTG. Prior to imaging, *E. coli* cells were grown in 1 mM IPTG for one hour then spotted on agar pads containing or not 10μg/ml cephalexin and incubated for 1 hour. Cell fluorescence profiles were obtained with the “plot profile” function of FIJI [49]. FrzCD clusters numbers and distances, nucleoid areas and cell areas were automatically determined and verified manually with the “MicrobeJ” Fiji/ImageJ plugin created by A. Ducret, Brun Lab (http://www.indiana.edu/~microbej/index.html). All data plots and statistical tests were obtained with the R software (https://www.r-project.org/).

To study the dynamic of FrzCD-GFP clusters we automatically tracked clusters (imaged every second) by MicrobeJ and recorded parameters such as the mean square displacement (MSD), the confinement radius and the fluorescence intensity along the time.

For manual image analysis we chose a sample size that allowed an error on the mean (sem) lower than 10%. For results generated by automated analyses, we analyzed all the available samples. We chose all cells where clusters and nucleoids were tractable by our image analysis tools (Fiji and MicrobeJ). For the cluster dynamic determinations we analyzed all clusters that were tractable for at least 5 consecutive frames.

### Reversal frequencies

These assays were performed as previously described [25] by using homemade PDMS glass microfluidic chambers [50] treated with 0.015% carboxymethylcellulose after extensive washing of the glass slide with water. For each experiment, 1mL of a CYE grown culture of OD = 0.5 was injected directly into the chamber and the cells were allowed to settle for 5 min. Motility was assayed after the chamber was washed with TPM 1mM CaCl buffer. For IAA injections, IAA solutions made in TPM 1mM CaCl buffer at appropriate concentrations were injected directly into the channels and motility was assayed directly under the microscope. Time-lapse movies of strains carrying a FrzS-YFP fusion were shot for 20 minutes with frames captured every 15 seconds.

To discriminate bona fide reversal events from stick-slip motions [25], the fluorescence intensity of FrzS-YFP was measured at cell poles over time. In fact, this protein has been shown to switch from the leading cell pole to the lagging pole when *M. xanthus* cells reverse their movement direction [51]. For each cell that was tracked, the fluorescence intensity and reversal profiles were correlated to distinguish bona fide reversals from stick-slip events. About one hundred cells for the wild type and *frzCD*^*Δ1-130*^ strains and fifty for the *ΔfrzCD* strain were analyzed (refer to Supplementary Table 3 for the exact number of cells analyzed for each strain and IAA dose). The number of reversals was plotted against time. The best fits for the reversal frequencies values at the different IAA doses were obtained with the following Hill equation:

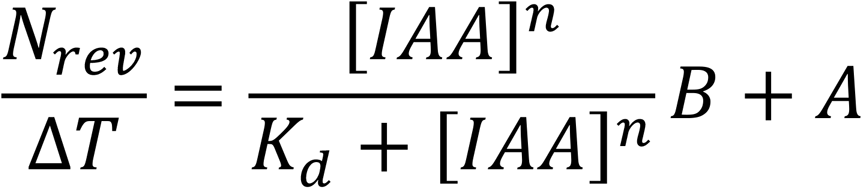

where the *N*_*rev*_/*ΔT* is the number of reversal events per hour; *K*_*d*_ is the apparent affinity constant; *[IAA]* is the IAA dose; *B* is the plateau; *A* is the basal reversal frequency and *n* is the Hill coefficient describing cooperativity. Reversal frequency values for each IAA dose and each strain are the results of two independent biological triplicates.

We chose a sample size that allowed an error on the mean (sem) lower than 10%. We used all cells that moved, remained isolated for at least 20 consecutive frames and where FrzS-YFP foci were detectable by FIJI.

## Acknowledgments

We would like to thank Dr. Romain Mercier and Dr. Thierry Doan for their critical reading of the manuscript and discussions; Dr. Mireille Ansaldi for her advise on the EMSA assays; Hanna Bismuth for helping with the ChIP experiments; Pravin Dewangan and Dr. Radha Chauhan for use of the SEC-MALS facility at National Centre for Cell Science, Pune, India.

Research on chemotaxis in our laboratory is funded by the Agence National de la Recherche Jeune Chercheur-Jeune Chercheuse (ANR-14-CE11-0023-01) to E.M.M and the “Fondation Amidex” award to EM and TM. YM, PJJ and PG acknowledge fellowships from INSPIRE, Department of Science and Technology (DST), Govt. of India and the work in the lab at IISER Pune is supported by Innovative Young Biotechnologist Award (IYBA from Department of Biotechnology, Govt. of India), Indian National Science Academy (INSA), and Science and Engineering Research Board (SERB), DST.

## Supplementary figure legend

**Figure S1.**
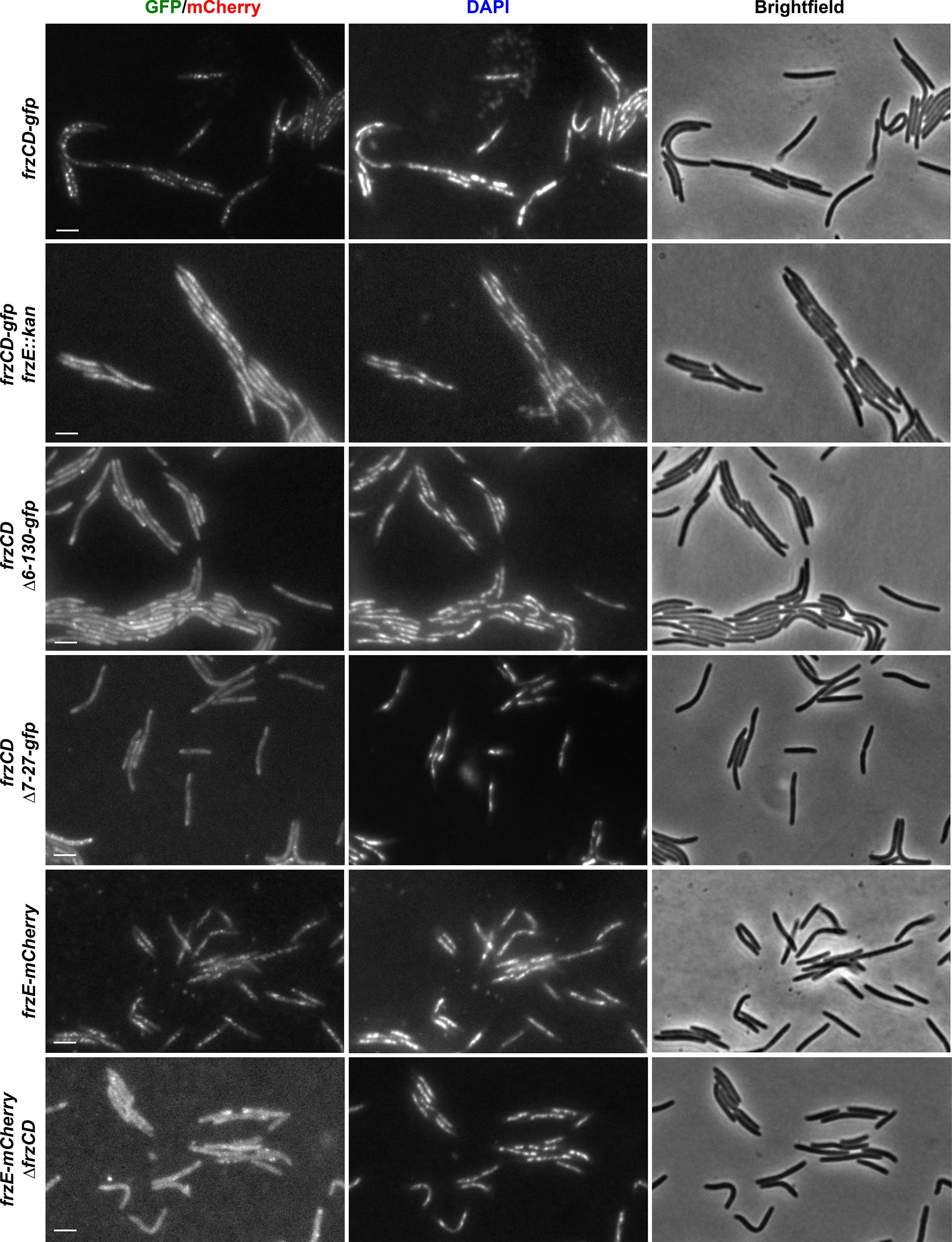
Representative micrographs of *M. xanthus* cells carrying a GFP or a mCherry fusion and stained with the DNA DAPI stain. The genetic backgrounds of the *M. xanthus* strains are indicated on the left. Scale bars correspond to 1 μm.

**Figure S2.**
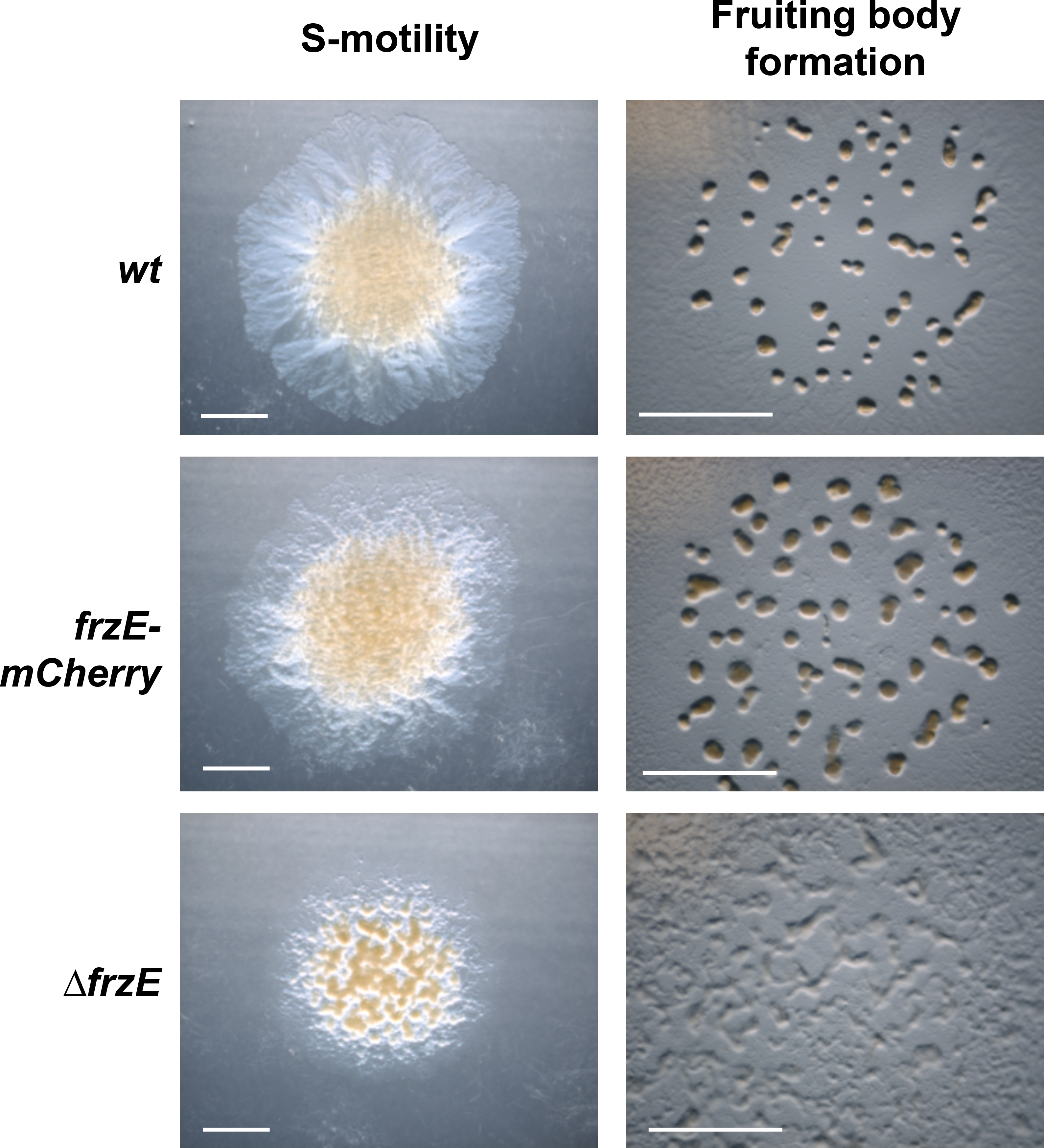
*frzE-mcherry* cells can swarm and form fruiting bodies like wild type. Scale bars correspond to 0,5 cm.

**Figure S3.**
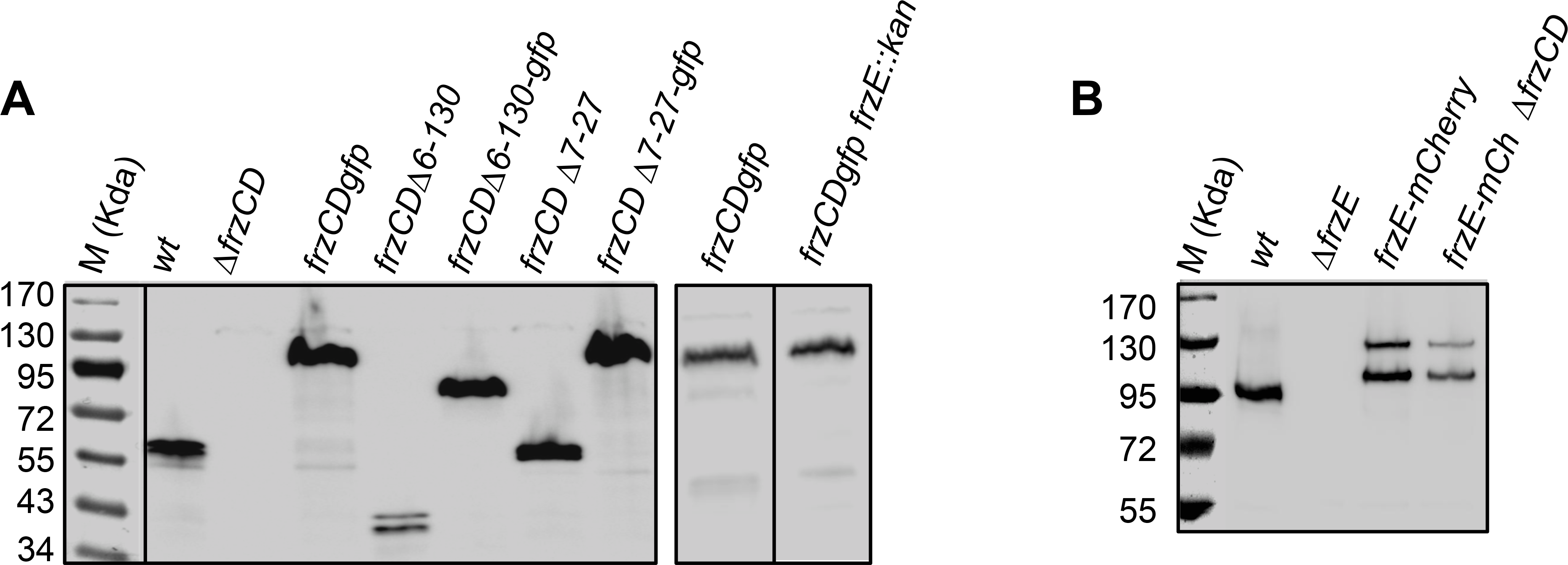
FrzCD and FrzE are stable in different *M. xanthus* mutants. Western blot with anti-FrzCD **(A)** or anti-FrzE antibodies **(B)** on the cell extracts of the indicated *M. xanthus* strains. Scale bars correspond to 1μm. Black lines are used to indicate that two lanes from the same gel where separated by other lanes in the original western blot.

**Figure S4.**
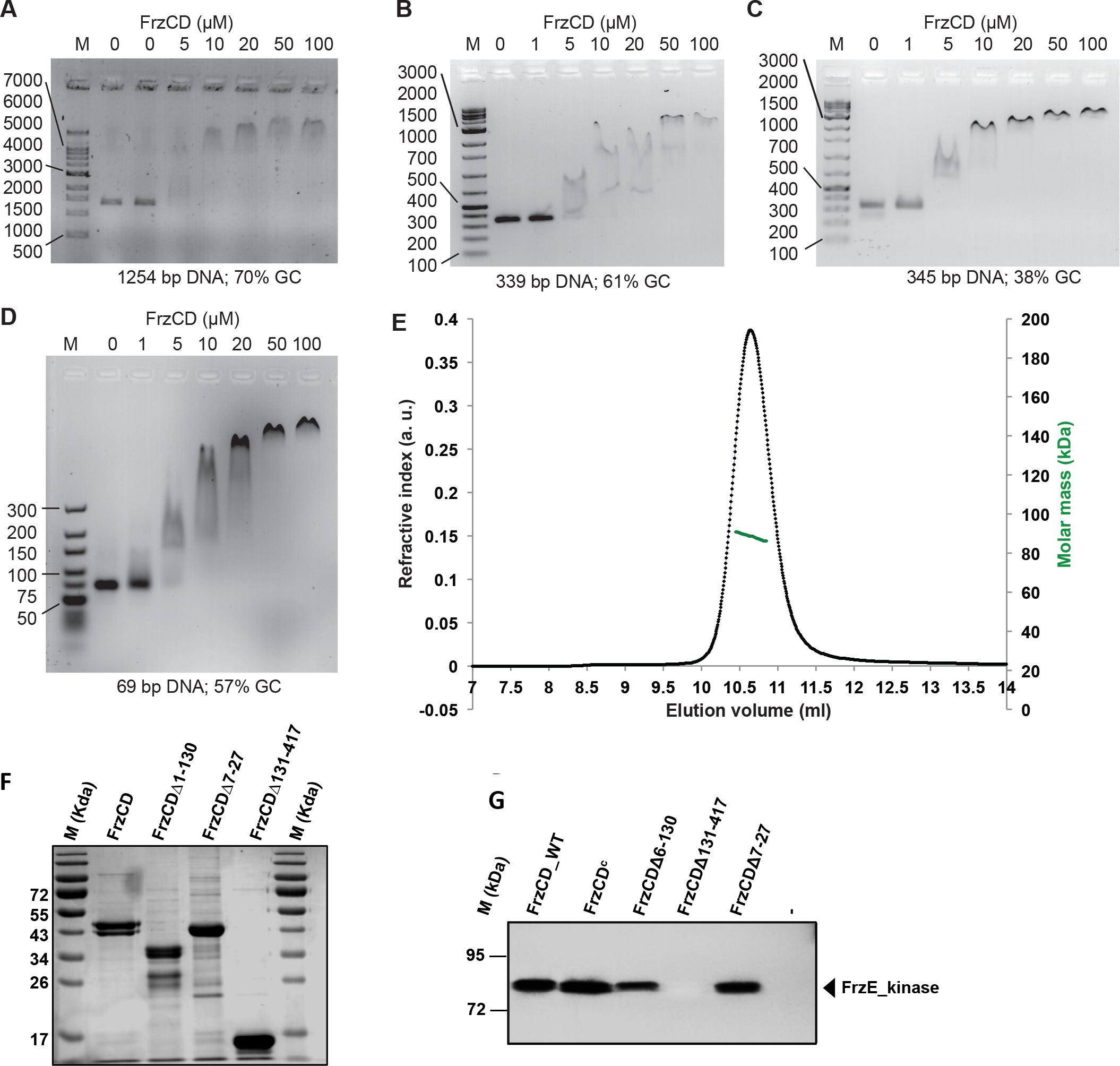
FrzCD binds DNA fragments of different lengths, forms a dimer and promotes FrzE phosphorylation. **(A- D)** Electrophoretic mobility shift assay (EMSA) on agarose gel stained with ethidium bromide and developed at the UV light. DNA fragments of different lengths and GC contents (Table S4) were incubated with increasing concentrations of FrzCD-(His_6_) as indicated, including **(A)** 4 nM of Oligo1300 (1% agarose gel), **(B)** 40 nM of 0ligo340-1 (38% GC content; 1.5% agarose gel), **(C)** 40 nM of Oligo340-2 (61% GC content; 1.5% agarose gel), and **(D)** 300 nM of Oligo70 (2.5% agarose gel). **(E)** SEC-MALS analysis showing the elution profile (refractive index; black; left y-axis) and the estimated molar masses (green; right y-axis) of the FrzCD eluted protein. **(F)** SDS page of the indicated proteins purified from *E. coli* and used for the different experiments shown in Figure 3. **(G)** Kinetics of the FrzE kinase domain (FrzE^CheA^) auto-phosphorylation were tested *in vitro* by incubation of FrzE^CheA^ in the presence of FrzA, the indicated different form of FrzCD and ATPγP^33^ as a phosphate donor.

**Figure S5.**
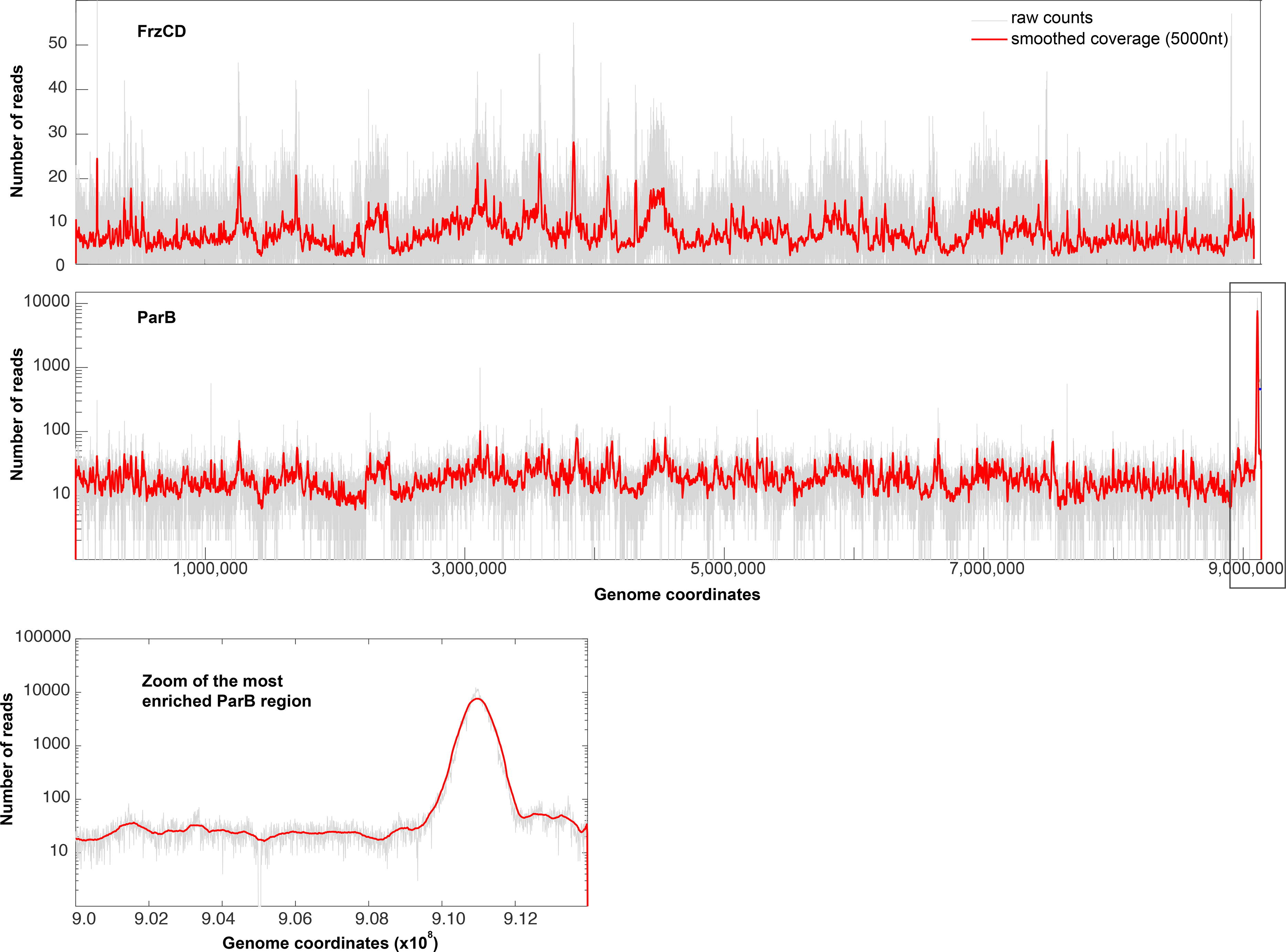
FrzCD-GFP binds the nucleoid in a DNA-sequence independent manner. A library of DNA fragments was obtained by ChIP experiments on *frzCD-gfp* and *parB-yfp* strains, using GFP polyclonal antibodies. The figure shows the results obtained by the deep sequencing of the DNA libraries. Only for *parB-yfp*, we observed an enrichment corresponding a to the nucleoid region containing *parS* (rectangle in the middle panel and last panel) [30] (9,109 to 9,110 Kb). Note that while the number of reads relative to ParB are represented with a logarithmic scale, for FrzCD we used a regular scale.

**Figure S6.**
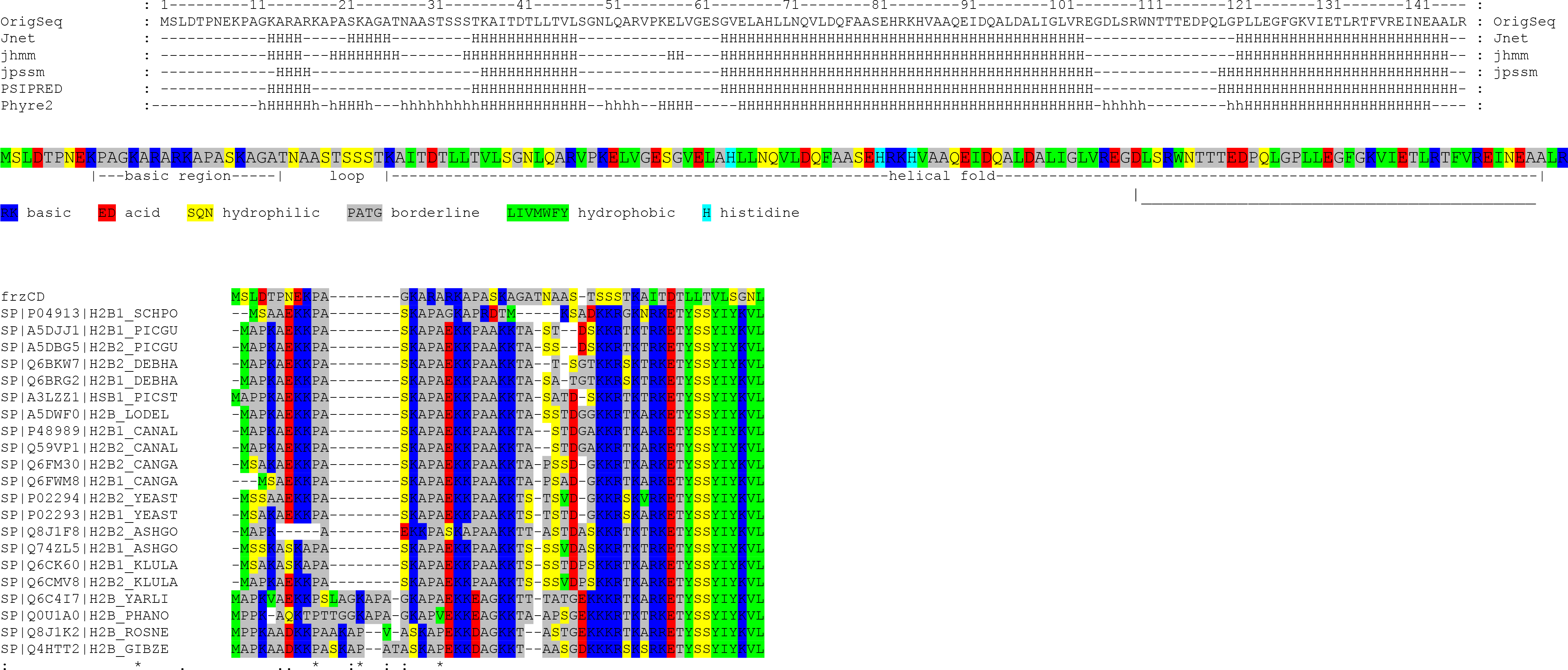
The FrzCD N-terminal tail has the same properties of that of eukaryotic histones. **(A)** Prediction of the FrzCD N-terminal secondary structures. The nature of each amino acid is also indicated through color codes. **(B)** FrzCD first 50 amino acid alignment with the N-terminal tail of Histones 2B. The alignment was obtained by Clustal Omega. Dots indicate similarities and stars identities.

**Figure S7.**
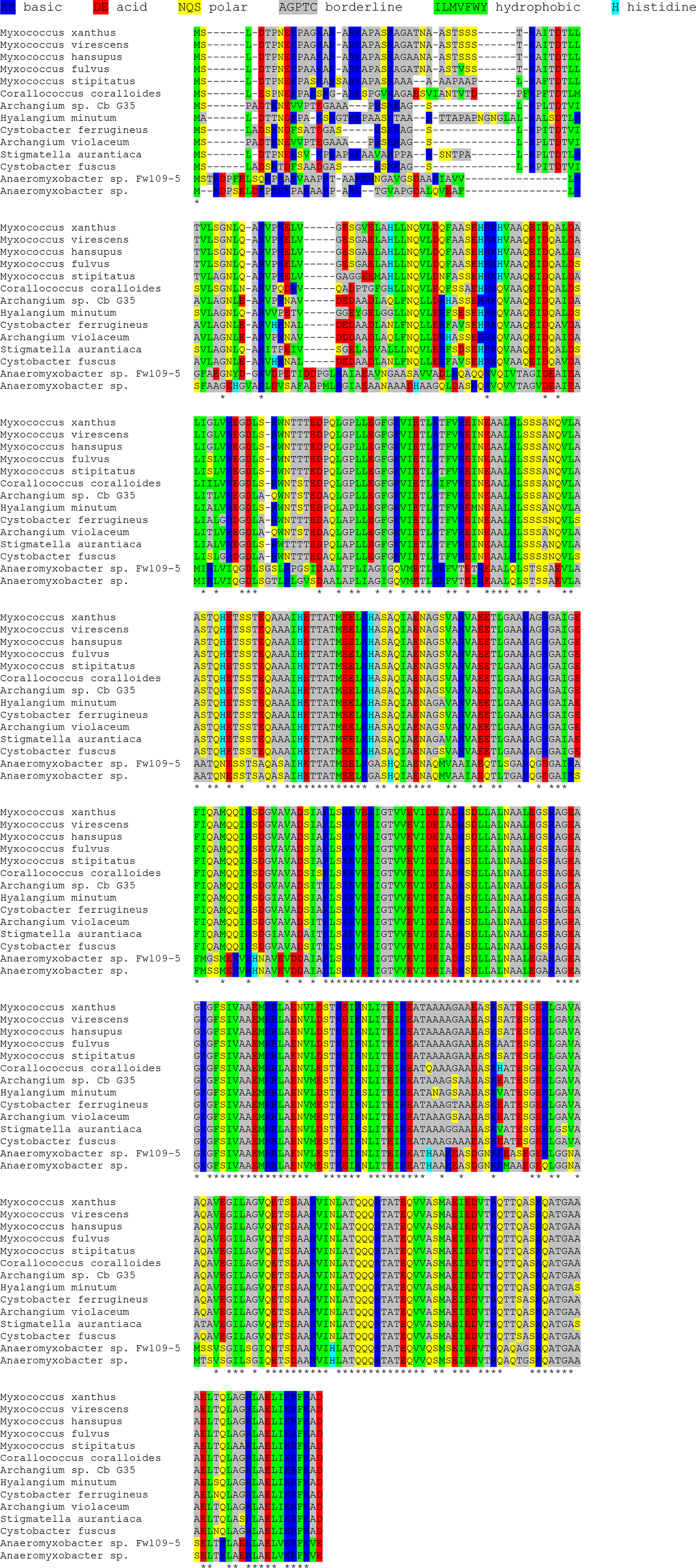
The FrzCD N-terminus from different species show similar amounts of positively charged amino acids. The alignment of FrzCD sequences from the indicated species was obtained by Clustal Omega. Stars indicate identities.

**Table S1:** Strains used in this study

| Strains | Genotyp | Deletion | Source |
| --- | --- | --- | --- |
| DZ2 | *wt* |  | Zusman et al., 1982 |
| DZ4620 | *frzCD-gfp* |  | Mauriello et al., 2009 |
| DZ4480 | $\Delta frzCD$ | Codons 6-393 | Bustamante et al., 2004 |
| DZ4485 | *frzCD*<sup>$\Delta 6-130$</sup> |  | Mauriello et al., 2009 |
| DZ4743 | *frzCD*<sup>$\Delta 6-130$</sup>-*gfp* |  | Mauriello et al., 2009 |
| EM231 | *frzCD*<sub>E202A-E203A::gfp</sub> |  | Mauriello et al., 2009 |
| EM228 | *frzCD*<sub>E168A-G169A::gfp</sub> |  | Mauriello et al., 2009 |
| EM434 | *frzE-mCherry* |  | This study |
| EM506 | *frzE-mCherry* $\Delta frzCD$ | | This study |
| EM516 | *frzCD-gfp frzE::kan* | Codons 171-438 | This study |
| EM531 | *difA-gfp* $\Delta parB/P_{cuoA}$-*parB* | | This study |
| EM532 | *frzCD-gfp* $\Delta parB/P_{cuoA}$-*parB* | | This study |
| EM533 | *frzE-mCherry* $\Delta parB/P_{cuoA}$-*parB* | | This study |
| EM543 | *frzCD*<sup>$\Delta 7-27$</sup> | Codons 7-27 | This study |
| EM550 | *frzCD*<sup>$\Delta 7-27$</sup>-*gfp* | Codons 7-27 | This study |
| TM26 | *frzS-yfp* |  | Guzzo et al., 2015 |
| EM622 | *frzCD*<sup>$\Delta 6-130$</sup> *frzS-yfp* |  | This study |
| EM623 | $\Delta frzCD$ *frzS-yfp* | | This study |

**Table S2:**
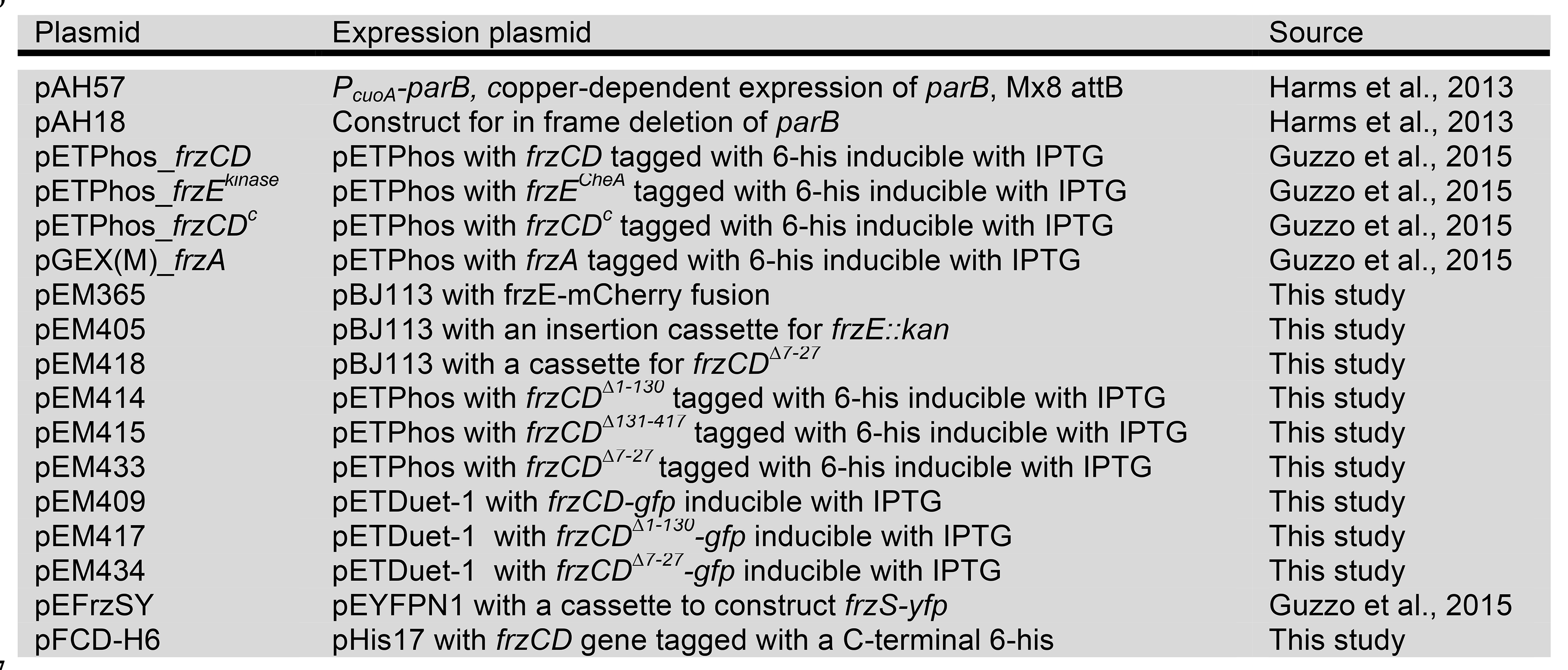
Plasmids used in this study

**Table S3:** Number of cells analyzed for Figure 5A

| %IAA | 0 | 0.005 | 0.01 | 0.03 | 0.05 | 0.1 | 0.3 |
| --- | --- | --- | --- | --- | --- | --- | --- |
| Strain |  |  |  |  |  |  |  |
| *frzS-yfp* | 101 | 113 | 77 | 114 | 64 | 47 | 170 |
| *frzCD*<sup>$\Delta 6-130$</sup> *frzS-yfp* | 98 | 103 | 75 | 128 | 112 | 139 | 109 |
| $\Delta frzCD$ *frzS-yfp* | 44 | ND | ND | ND | 48 | ND | 47 |

**Table S4:**
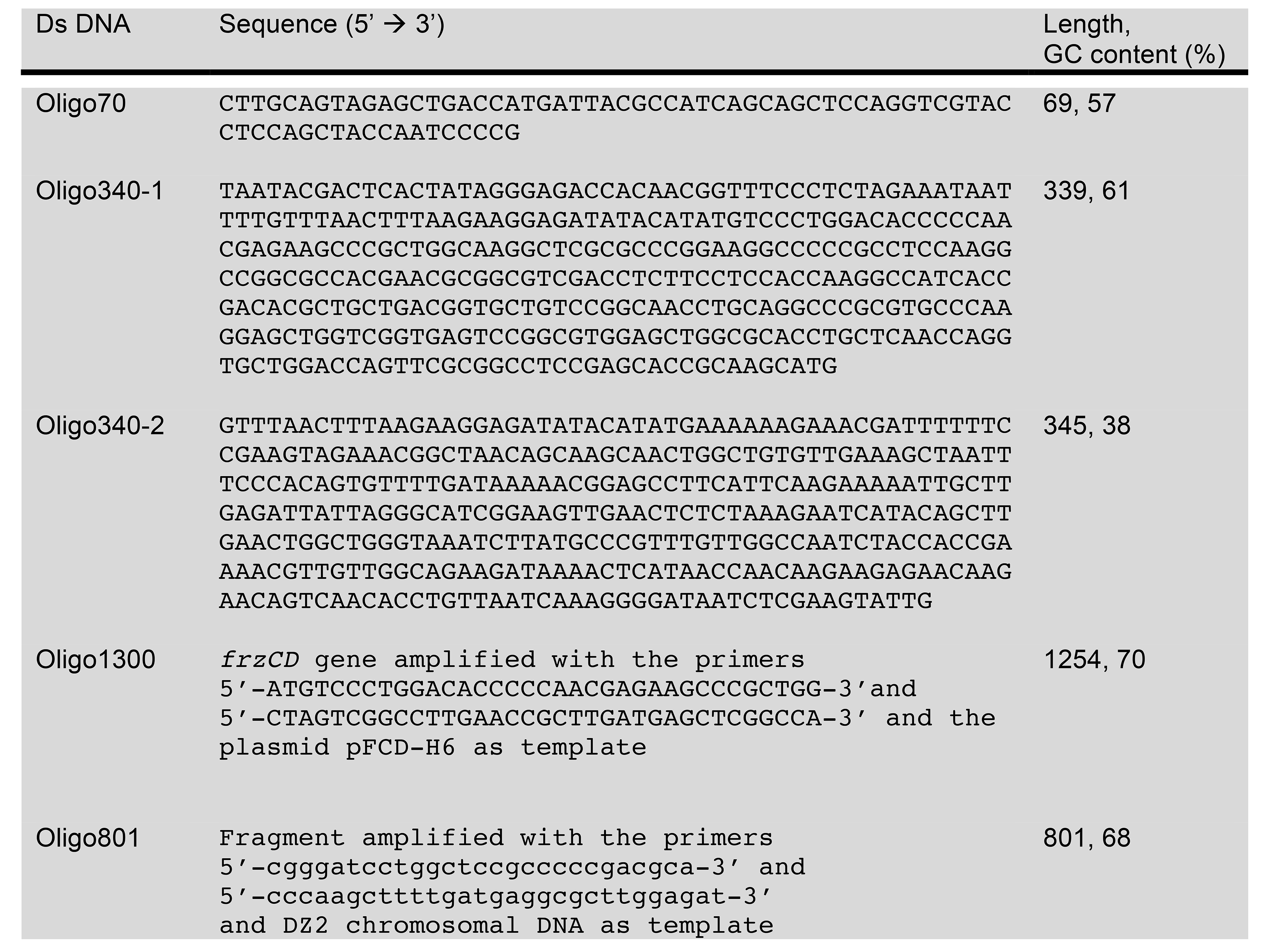
DNA sequences used for EMSA in Figure 3 and Supplementary figure 4

